# Clastogenesis by nucleotide lesions requires the completion of two cell cycles

**DOI:** 10.64898/2025.12.16.694654

**Authors:** Jacob G. Jansen, Piya Temviriyanukul, Daniel C. de Groot, Karoly Szuhai, Sandrine van Hees-Stuivenberg, Anastasia Tsaalbi-Shtylik, Heinz Jacobs, Niels De Wind

## Abstract

Damaged DNA nucleotides can trigger genome rearrangements through clastogenesis, a process driven by erroneous repair of double-strand breaks (DSBs) and associated with cancer development. While DSBs are known to arise from endonuclease activity at stalled replication forks, the clastogenic potential of such DSBs has remained uncertain. Here, we identify a previously unrecognized mechanism of clastogenesis using wild-type, nucleotide excision repair (NER)-deficient and translesion synthesis (TLS)-deficient cells, combined with advanced cytogenetic analyses. We demonstrate that, single-stranded DNA (ssDNA) tracts harboring unrepaired lesions rather than DSBs at collapsed replication forks can persist through mitosis. Only during the subsequent S phase, these tracts are converted into a new class of, highly clastogenic, DSBs. Consistent with a role of this mechanism in carcinogenesis, prostate cancers exhibiting extensive genomic rearrangements frequently harbor somatic defects in NER or in error-free homologous recombination-mediated DSB repair. These findings provide critical mechanistic insight and highlight potential implications for routine clastogenicity testing.

**Graphical abstract:** Nucleotide lesions (light blue triangle) can trigger double-strand breaks (DSBs) through endonucleolytic cleavage at stalled or reversed replication forks. Traditionally, these DSBs were assumed to drive genome rearrangements, a process termed clastogenesis. Here we describe a distinct, delayed, mechanism of clastogenesis. Thus, unreplicated nucleotide lesions within single-stranded (ss) DNA regions persist through mitosis into the next cell cycle. During the subsequent S phase, these ssDNA tracts collapse into DSBs, presumably via replication runoff. These delayed DSBs then promote extensive genomic reshuffling. Supporting this model, prostate cancers with high levels of genomic rearrangements are frequently associated with somatic defects in nucleotide excision repair (NER)—a pathway that normally prevents lesion-induced clastogenesis.

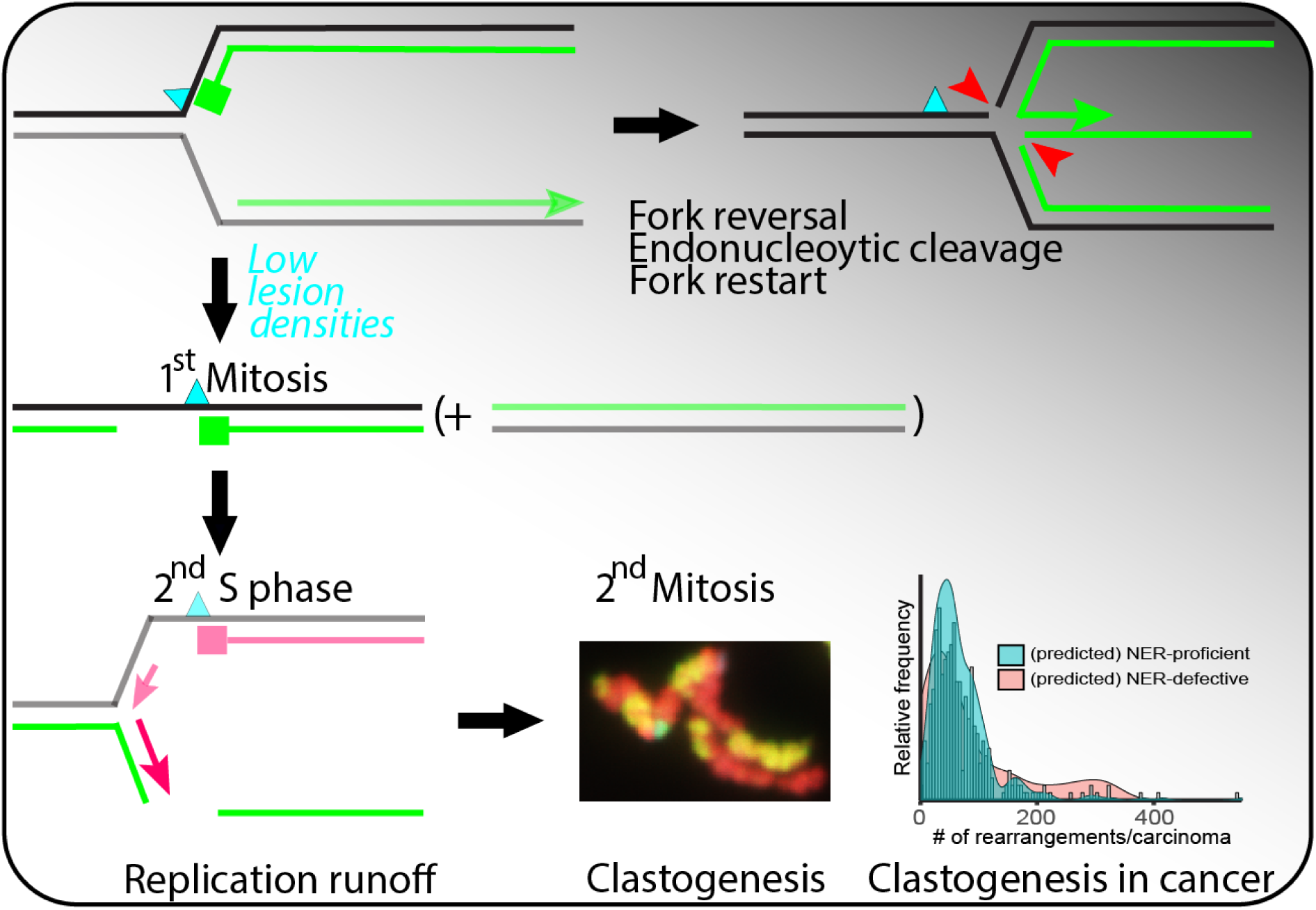

## Introduction

Prostate and other cancers frequently arise in the context of chronic low-level exposure to endogenous and exogenous DNA-damaging agents ^1, 2^. These malignancies are characterized by both nucleotide substitution mutations and extensive genome rearrangements ^3, 4^. While mutational signature analysis has provided critical insights into the relationship between nucleotide lesions and nucleotide substitutions ^5, 6^, the contribution of nucleotide lesions to clastogenesis—the process driving genome rearrangements—remains poorly defined.

Nucleotide lesions that evade removal by nucleotide excision repair (NER) stall replication forks, inducing replication stress ^7^. Fork restart beyond the lesion or convergence from adjacent replicons generates post-replicative single-stranded DNA (ssDNA) tracts (Fig. 1a; ^8, 9, 10^). These tracts are stabilized by replication protein A (Rpa) ^11^, which simultaneously induces Atr-dependent DNA damage responses, including checkpoint signaling, repair, senescence, or apoptosis ^12, 13^. Such lesion-containing ssDNA tracts can be filled by DNA translesion synthesis (TLS). Thereby, TLS enables replication completion, which alleviates replication stress albeit at the cost of frequent misincorporations that generate nucleotide substitutions ^10, 14^. Alternatively, a replication fork stalled at a lesion may reverse to form a “chicken foot” structure, which allows the undamaged sister chromatid to serve as a template for replication (Fig. 1a). Cleavage of chicken feet by endonucleases such as Mus81/Eme1 can produce double-strand breaks (DSBs), called collapse of replication forks. Such DSBs enable fork remodeling and restart via homologous recombination (HR) ^10, 11^. Additional sources of DSBs include ionizing radiation, opposing NER-induced ssDNA tracts ^15^, replication runoff at excision repair-induced excision tracts ^16, 17^, breakage of anaphase bridges during mitosis ^18, 19^, and incomplete replication of chromosomes or micronuclear fragments ^20^. It is evident that clastogenesis, a hallmark of carcinogenesis ^3, 4^, requires the induction and mis-repair of DSBs ^11, 21^. Nevertheless, the causal involvement of DSBs at collapsed replication forks in clastogenesis has not been proven, particularly at physiologically relevant lesion densities.

**Figure 1:**
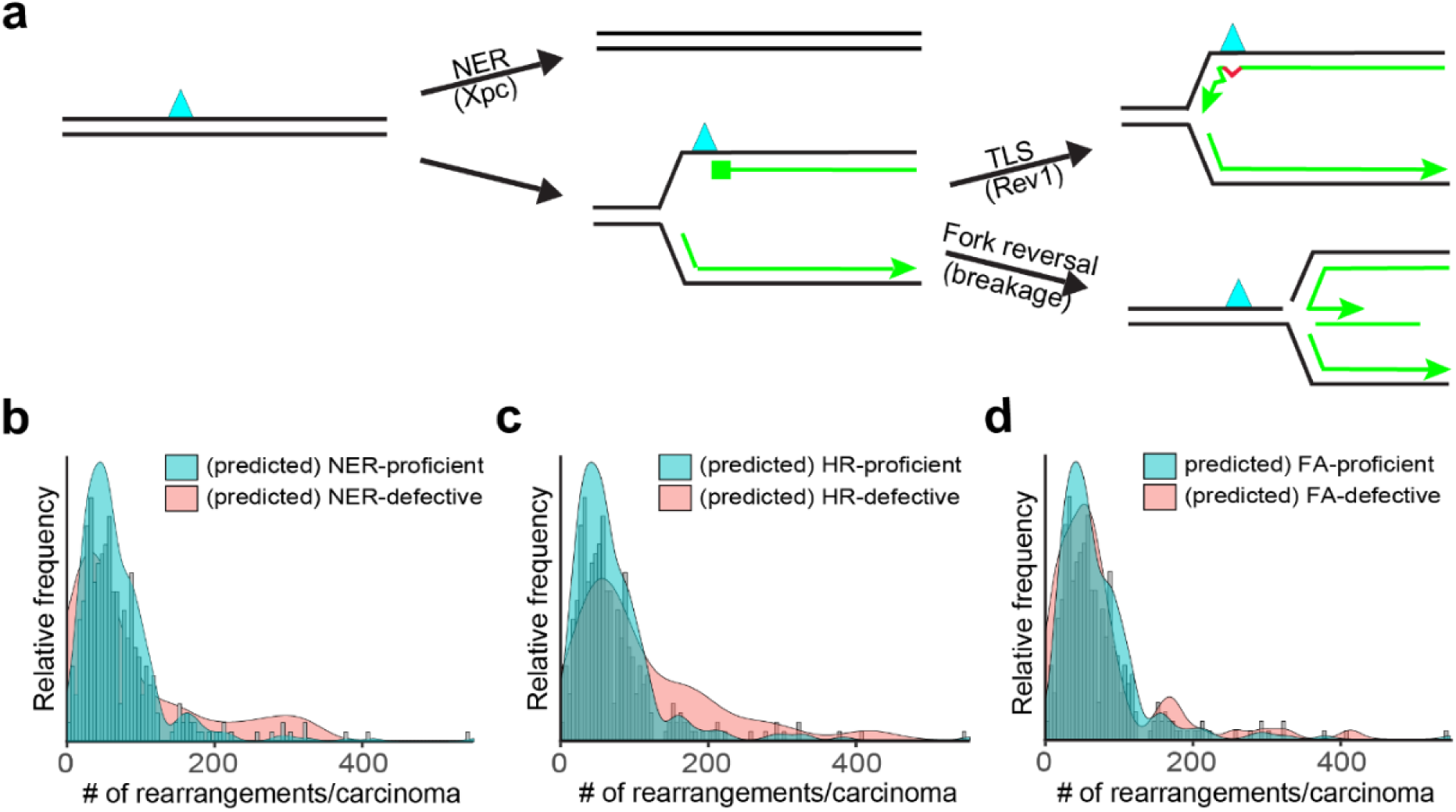
Enrichment of NER and HR defects in highly genome-unstable prostate carcinomas. a) NER and TLS, respectively, repair and replicate helix-distorting nucleotide lesions (blue triangle). Red: TLS-induced misincorporation. Alternatively, the fork can reverse into a “chicken foot”, which not only enables the leading strand to use the undamaged sister as a template for replication but also is also prone to endonucleolytic cleavage, resulting in a dsDNA break. b) Prevalence of (predicted) benign and deleterious mutations in NER-associated genes (density plots) as a function of the number of genome rearrangements per carcinoma (bars). c) Prevalence of (predicted) benign and deleterious mutations in HR-associated genes (density curves) as a function of the number of genome rearrangements per carcinoma (bars). d) Prevalence of (predicted) benign and deleterious mutations in FA-associated genes (density curves) as a function of the number of genome rearrangements per carcinoma (bars).

To elucidate mechanisms of clastogenic DSB formation we exposed isogenic mouse embryonic fibroblast (MEF) lines, either wild type or deficient in NER or TLS, during S phase to doses of UVC light, that allow cell cycle progression. Using specialized cytogenetic approaches, we tracked photolesions and genomes across consecutive cell cycles. Our findings reveal that Rpa-coated ssDNA tracts containing unreplicated photolesions persist through mitosis, collapsing into DSBs only during the subsequent S phase. These delayed DSBs are highly clastogenic, mediating extensive genome rearrangements. We propose that this mechanism contributes to carcinogenesis associated with chronic low-level DNA damage. Supporting this hypothesis, prostate carcinomas with many genomic rearrangements are enriched for predicted deleterious mutations in NER and HR genes.

## Results

### Somatic mutations in NER and HR-associated genes in highly genome-rearranged prostate carcinomas

Prostate cancer is linked to chronic, low-level exposure to endogenous and exogenous sources of DNA-damage ^1, 2^. A subset of prostate and other cancers exhibits extensive genome-wide rearrangements ^4, 22^. Assuming that these rearrangements are driven by DNA lesion–induced DSBs, we hypothesized that the most highly rearranged prostate carcinomas would be enriched for somatic defects in pathways that suppress the clastogenic potential of DNA lesions. These pathways include nucleotide excision repair (NER), interstrand crosslink repair via the Fanconi anemia (FA) pathway—which relies on TLS proteins ^23^—and homologous recombination (HR), as loss of HR-mediated error-free DSB repair is expected to increase reliance on error-prone, recombinogenic, DSB repair mechanisms ^24, 25, 26^.

To test this hypothesis, we analyzed whole-genome sequences of prostate cancers for predicted deleterious somatic mutations in genes associated with NER, FA, and HR (Supplementary Table 1). Among tumors with the highest numbers (>200) of genome rearrangements, we observed significant enrichment of mutations in NER genes (4.6-fold, p = 0.02; Fig. 1b, Supplementary Table 2) and HR genes (3.9-fold, p = 0.03; Fig. 1c, Supplementary Table 2). In contrast, mutations in FA genes were not significantly associated with extreme genome instability (p = 0.33; Fig. 1d, Supplementary Table 2), consistent with recent evidence that FA activity is required for the induction of genome rearrangements ^27^. Moreover, TLS contributes to survival under replication stress induced by oncogene activation ^28, 29, 30, 31^ and therefore somatic TLS defects may not be compatible with cancer development. Collectively, these findings support a model in which clastogenic nucleotide lesions and error-prone DSB repair contribute to the genesis of highly rearranged prostate cancers.

### *Xpc* and *Rev1Xpc* MEFs are suitable models to study responses to low UVC fluencies

Our observation that highly genome-rearranged prostate carcinomas frequently harbor NER defects—suggesting a link between persisting nucleotide lesions and clastogenesis—motivated the use of NER-deficient (*Xpc^⁻/⁻^*) mouse embryonic fibroblasts (MEFs) as a sensitive model to dissect the mechanistic basis of clastogenesis. Moreover, Xpc deficiency eliminates NER-induced excision tracts as a potential source of clastogenic DSBs, thereby avoiding confounding effects ^15^.

We employed UVC irradiation as a model clastogen for proliferating cells ^32^, which generates severely distorting pyrimidine-pyrimidone (6-4) photoproducts [(6-4)PP] and moderately distorting cyclobutane pyrimidine dimers (CPDs) at an approximate ratio of 1:3 ^33, 34^. Among these, (6-4)PP lesions strongly arrest replication forks, leading to single-stranded DNA (ssDNA) tracts that activate ATR signaling ^35^. Rev1-dependent TLS provides one mechanism to tolerate (6-4)PP ^36, 37^. To specifically study clastogenesis by unreplicated (6-4)PP lesions we therefore additionally disrupted *Rev1*.

As anticipated, *Xpc*-deficient MEFs exhibited moderate sensitivity to UVC, compared with wild-type cells, whereas *Rev1Xpc* double-deficient MEFs displayed synergistically increased sensitivity (Supplementary Fig. 1). These findings are consistent with the roles of Xpc and Rev1 in NER and TLS, respectively of (6-4)PP photolesions ^36^.

### Low UVC fluencies rarely induce micronuclei and genome rearrangements, shortly after UVC exposure

To assess whether low-dose UVC exposure induces DSBs, we monitored phosphorylation of Kap1 (pKap1), a substrate of the Ataxia telangiectasia-mutated (ATM) kinase activated by DSBs ^38^. In *Xpc* and *Rev1Xpc* MEFs replicating at the time of equitoxic low-dose UVC exposure we observed a marked increase in pKap1 within 4 hours (Fig. 2a). This indicates rapid DSB formation, presumably resulting from the collapse of stalled replication forks. To determine whether these early DSBs persist through mitosis, we employed the cytokinesis-block micronucleus assay ^39^. Thus, immediately after UVC exposure, cells were treated with Cytochalasin B to block cytokinesis, enabling the facile identification of binucleated post-mitotic cells ^40^. At 24 hours post mock treatment, no micronuclei were detected in binucleated *Rev1Xpc* or *Xpc* cells, confirming that Cytochalasin B itself is not clastogenic. Unexpectedly, micronuclei were also not observed at 24 hours following low-dose UVC exposure (Fig. 2b), suggesting that UVC-induced early DSBs are either repaired rapidly or not transferred through mitosis. We then investigated early DSBs persist to mitosis by using the neutral single-cell electrophoresis (“comet”) assay of *Rev1Xpc* MEFs, cultured in the presence of nocodazole. No comets indicative of DSBs were detected at 16 hours post-UVC treatment, suggesting the efficient repair of these DSBs prior to mitosis (Fig. 2c).

**Figure 2:**
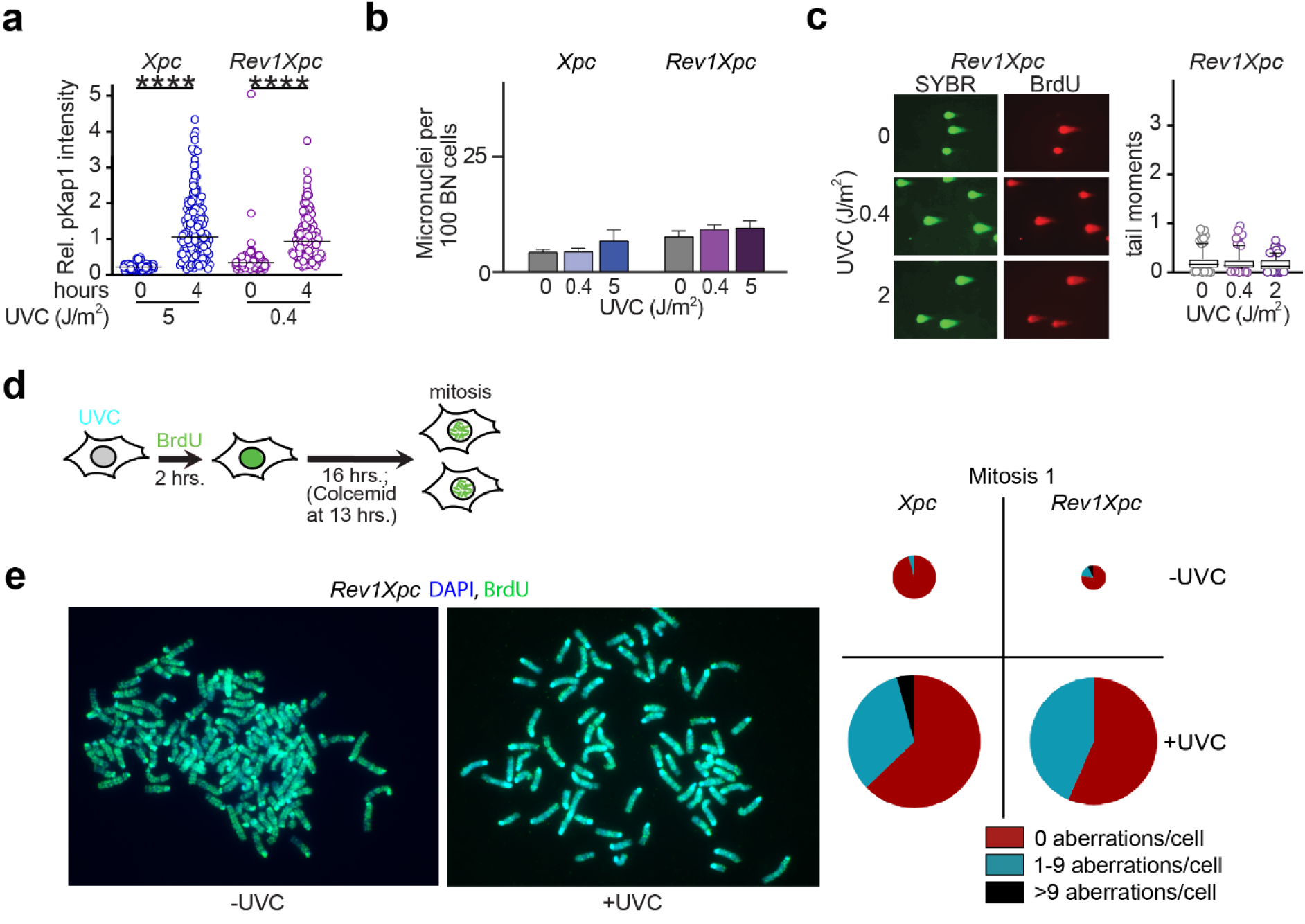
Low clastogenic activity of dsDNA breaks induced during the cell cycle of low dose UVC exposure. a) Quantification of immunostainings for pKap1 in EdU positive *Xpc* and *Rev1Xpc* cells at 0 and 4 hours after UVC exposure. Cells were pulse-labelled with the replication marker EdU, immediately following exposure. Cells were arrested at mitosis using Nocodazole. Error bars: SD b) Frequencies of micronuclei in in post-mitotic (binucleated, BN) *Xpc* and *Rev1Xpc* cells, 24 hours after UVC exposure. Cytochalasin B was added immediately after exposure. c) Left panel: single-cell (comet) gel electrophoresis at neutral pH to detect dsDNA, at 16 hours after low-dose UVC exposure of *Rev1Xpc* cells during S phase (BrdU^+^ cells). Immediately following exposure, Nocodazole was added to arrest cells at mitosis. SYBR green was used to visualize DNA. Right panel: quantification of tail moments of BrdU^+^ cells from three independent comet experiments. The box-and-whiskers plots represent 5 to 95 percentiles. d) Outline of the experiment to assess genome aberrations at mitosis of the first cell cycle after low-dose UVC exposure during S phase. e) Left panel: *Rev1Xpc* metaphase of the cell cycle of UVC exposure (0,4 J/m^2^) during S phase. Right panel: Quantification of genome aberrations in *Rev1Xpc* and *Xpc* metaphases of the cell cycle of low dose UVC exposure (0,4 and 5 J/m^2^, respectively) or mock treatment. The size of each circle represents the number of metaphases analyzed (between 13 and 70).

To directly assess the induction of clastogenesis by low-dose UVC light, we examined metaphase spreads from *Xpc* and *Rev1Xpc* MEFs, 16 hours after equitoxic low-dose UVC exposure or mock treatment during S phase (Fig. 2d). In mock-treated cells, only very few mitoses were present, likely because most cells had already progressed to the second cycle, as confirmed by fluorescence-activated cell sorting showing re-entry into G1 (Supplementary Fig. 2). UVC exposure did delay cell cycle progression, and therefore many metaphases were present at 16 hours following UVC exposure (Fig. 2e; Supplementary Fig. 2). However, only a minority of these metaphases exhibited UVC-induced aberrations (Fig. 2e). Collectively, these findings suggest that at the UVC doses tested, replication-associated DSBs are induced but that most of these are repaired prior to mitosis, precluding marked clastogenesis.

### Low UVC fluencies induce DSBs and genome rearrangements, but only with delayed kinetics

We hypothesized that clastogenicity of low-dose UVC might rather involve the delayed induction of recombinogenic DSBs. To test this, we again quantified micronucleus formation in *Xpc* and *Rev1Xpc* cells exposed to UVC, but now adding Cytochalasin B only after 24 hours. Indeed, at 48 hours post-exposure, a strong increase in micronuclei was seen in binucleated *Xpc* and *Rev1Xpc* MEFs (Fig. 3a), indicating a Rev1-independent mechanism of DSB induction that operates with delayed kinetics. Notably, *Rev1Xpc* cells exposed to a, for this genotype, highly cytotoxic dose (5 J/m²) displayed far fewer micronuclei than those exposed to 0.4 J/m², likely due to a strong cell cycle arrest precluding micronucleus formation.

**Figure 3:**
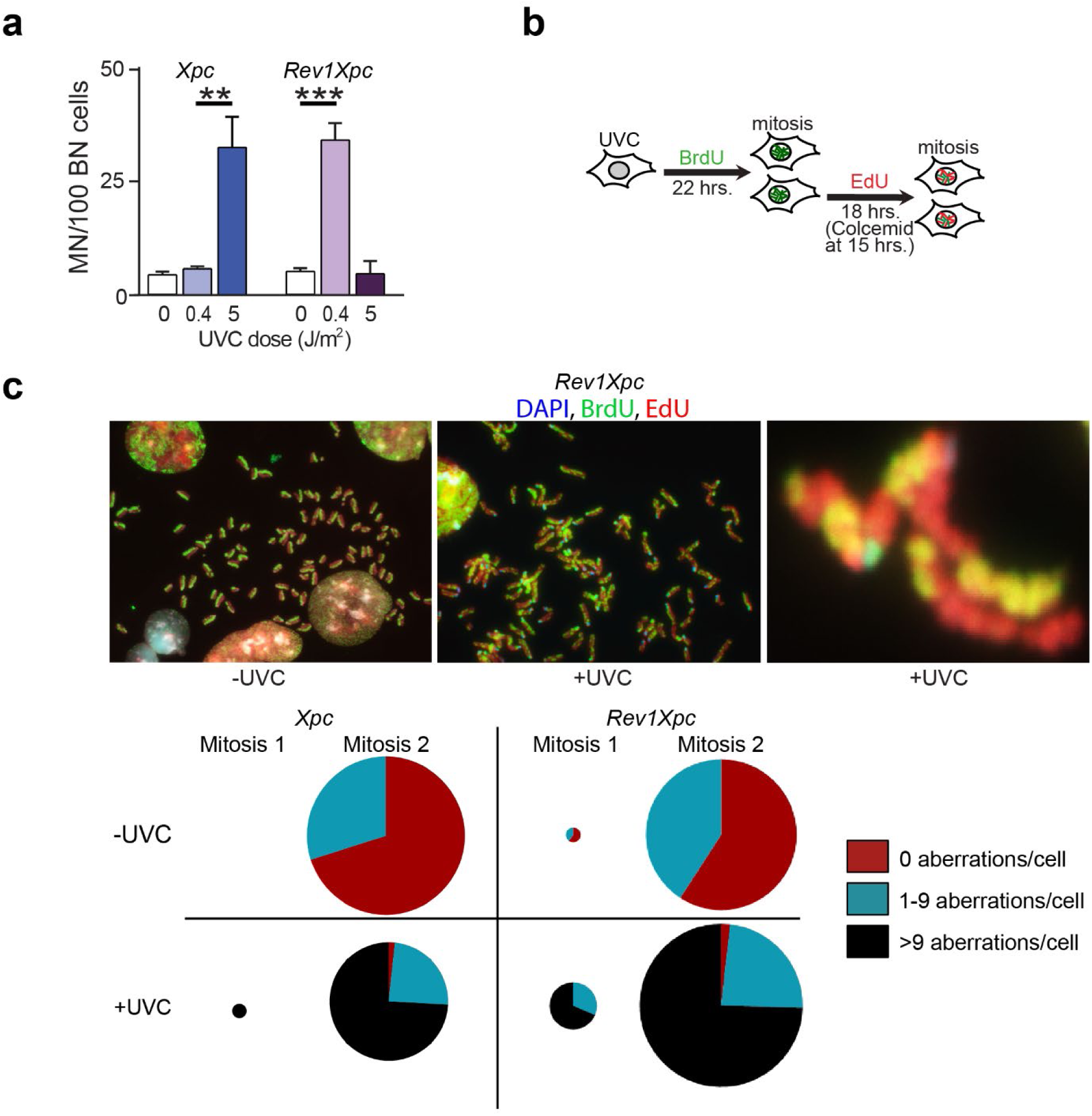
Clastogenesis by low lesion densities requires two cell cycles. a) Frequencies of micronuclei in in post-mitotic (binucleated, BN) *Xpc* and *Rev1Xpc* cells, 48 hours after UVC exposure. Cytochalasin B was added 24 hours after exposure. **: p<0.01, ***: p<0.001 (Student’s T test). b) Outline of the experiment to assess genome aberrations at mitosis of the second cell cycle after low dose UVC exposure. c) Top panel: metaphases of *Rev1Xpc* cells at mitosis of the second cell cycle after treatment with 0.4 J/m^2^ UVC before or during S phase. Ubiquitous sister chromatid exchanges and interchromosomal recombinations are apparent. Green: BrdU (first S phase), Red: EdU (second S phase), Blue: DAPI (centromere). Bottom panel: quantification of genome aberrations in metaphases of *Xpc* and *Rev1Xpc* cells after two cell cycles following low dose UV exposure or mock treatment during S phase. The size of each circle represents the number of metaphases analyzed (between 0 and 71).

We next examined whether the observed delayed DSB formation leads to clastogenesis during the second cell cycle. *Xpc* and *Rev1Xpc* MEFs were exposed to equitoxic low UVC doses and then differentially labeled during the first and second S phases using thymidine analogs 5-bromo-2’-deoxyuridine (BrdU) and 5-ethynyl-2’-deoxyuridine (EdU), respectively, followed by blockage at metaphase (Fig. 3b). At 40 hours post-exposure, most cells had reached the second mitosis, as indicated by BrdU staining restricted to one sister chromatid (representing the first S phase) while both chromatids were labeled with EdU (during the second S phase; Fig. 3c, Supplementary Fig. 3a). This result recapitulates the classical Meselson–Stahl demonstration of semi-conservative replication^41^. Strikingly, the majority of the UVC-exposed second-mitosis cells exhibited more than nine genomic rearrangements, a result consistent across *Xpc* and *Rev1Xpc* cells treated with equitoxic UVC doses, confirming the genotype-independence of this delayed clastogenesis mechanism (Fig. 3d). A minority of cells had remained in the first-cycle, at 40 hours post-exposure, as evidenced by symmetric BrdU staining of both chromatids (Fig. 3c). Also these lagging cells displayed extensive chromosomal rearrangements, possibly reflecting residual early DSBs (derived from collapsed replication forks) that have impeded progression to the second cell cycle.

### Delayed formation of dsDNA breaks following low UVC exposure

Given the extensive chromosomal aberrations observed in second-mitosis cells (Fig. 3c), we asked whether a novel, highly clastogenic, class of DSBs is induced during the second cell cycle. To address this, we analyzed binucleated cells originating from populations exposed to low-dose UVC during S phase (BrdU-labeled), followed by incubation with Cytochalasin B, to block cytokinesis while permitting cell cycle progression. Prior to fixation, EdU was added to label binucleated cells that had have entered the second S phase (BrdU⁺EdU⁺), enabling to distinguish these cells from those that have not yet entered the second S phase (BrdU⁺EdU⁻, Fig. 4a). Immunocytochemical staining revealed a significant increase in chromatin-bound Rad51—a marker of HR repair of DSBs ^42^—in cells during the second S phase (BrdU⁺EdU⁺), independent of genotype (Fig. 4b, 4c). This HR activity aligns with the frequent sister chromatid exchanges observed in second-mitosis metaphases, as evidenced by alternating BrdU staining between chromatids (Fig. 3c; Supplementary Fig. 3a). Consistently, levels of pKap1 were also elevated in second S-phase cells, in both genotypes exposed to equitoxic UVC doses (Fig. 4d, 4e). Together, these findings identify a previously unrecognized class of delayed DSBs that arise during the S phase of the cell cycle following that of UVC exposure. These delayed DSBs are highly clastogenic.

**Figure 4:**
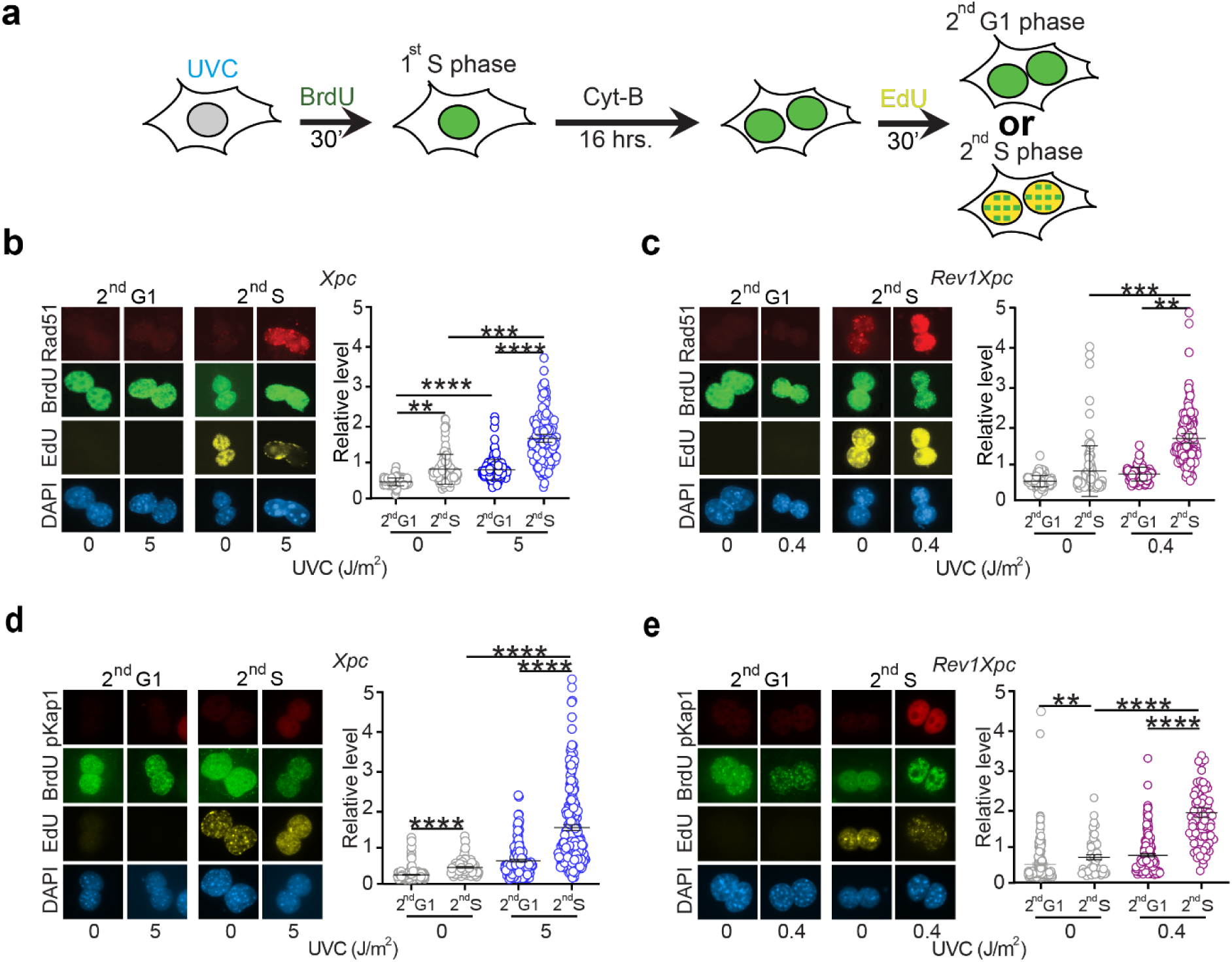
Delayed formation of recombinogenic dsDNA breaks during the S phase of the cell cycle following that of low dose UVC exposure. a) Outline of the experiments to analyze UVC-exposed cells at the second cycle after that of exposure to low-dose UVC light during S phase. b) Left panel: immunostainings for chromatin-bound Rad51 in binucleated (second-cycle) G1 and S phase *Xpc* cells. Right panel: quantification of chromatin-bound Rad51 in second cycle G1 and S phase *Xpc* cells. Error bars: S.D. **: p<0.01, ***: p<0.001, ****: p<0.0001 (Student’s T test). c) Left panel: immunostainings for chromatin-bound Rad51 in binucleated G1 and S phase *Rev1Xpc* cells. Right panel: quantification of chromatin-bound Rad51 in second cycle G1 and S phase *Rev1Xpc* cells. Error bars: S.D. **: p<0.01, ***: p<0.001 (Student’s T test). d) Left panel: immunostainings for phosphorylated Kap1 (pKap1) in binucleated G1 and S phase *Xpc* cells. Right panel: quantification of pKap1 in second cycle G1 and S phase *Xpc* cells. Error bars: S.D. ****: p<0.0001 (Student’s T test). e) Left panel: immunostainings for pKap1 in binucleated G1 and S phase *Rev1Xpc* cells. Right panel: quantification of immunostaining of pKap1 in binucleated G1 and S-phase *Rev1Xpc* cells. Error bars: S.D. **: p<0.01, ****: p<0.0001 (Student’s T test).

### Unreplicated (single-stranded) photolesions progress to mitosis, protected by Rpa

We next investigated whether this delayed DSB formation might result from the induction of ssDNA tracts immediately after exposure and that persist to the second cell cycle. To address this, we used *Rev1Xpc* cells that exhibit pronounced responses even at very low UVC doses. Comet electrophoresis under alkaline conditions at 16 hours post-exposure indeed revealed ssDNA discontinuities in mitosis-blocked *Rev1Xpc* cells that had been replicating during exposure (Fig. 5a). Of note, we detected no early DSBs at the same UVC dose and time point by comet electrophoresis under neutral conditions (Fig. 2c, see above). Immunocytochemical analysis of cells that had been in S phase during UVC exposure indeed showed chromatin-bound Rpa at 8 hours post-exposure (Fig. 5b), consistent with the presence of ssDNA tracts. Furthermore, staining with a monoclonal antibody specific for (6-4)PP embedded in ssDNA [ss(6-4)PP] ^36, 43^ confirmed that these ssDNA tracts represented unreplicated ss(6-4)PP (Fig. 5c). At this stage, Atr-phosphorylated Chk1 and γH2AX levels were also elevated (Fig. 5d; Supplementary Fig. 4), indicating activation of the DNA damage response, likely by ss(6-4)PP (or by the early transient DSBs). When replicating cells were exposed to UVC and arrested in mitosis, most phospho-Histone H3–positive (mitotic) cells retained ss(6-4)PP (Fig. 5e). This confirms that cells carrying unreplicated ss(6-4)PP enter mitosis, in agreement with the alkaline comet assays.

**Figure 5:**
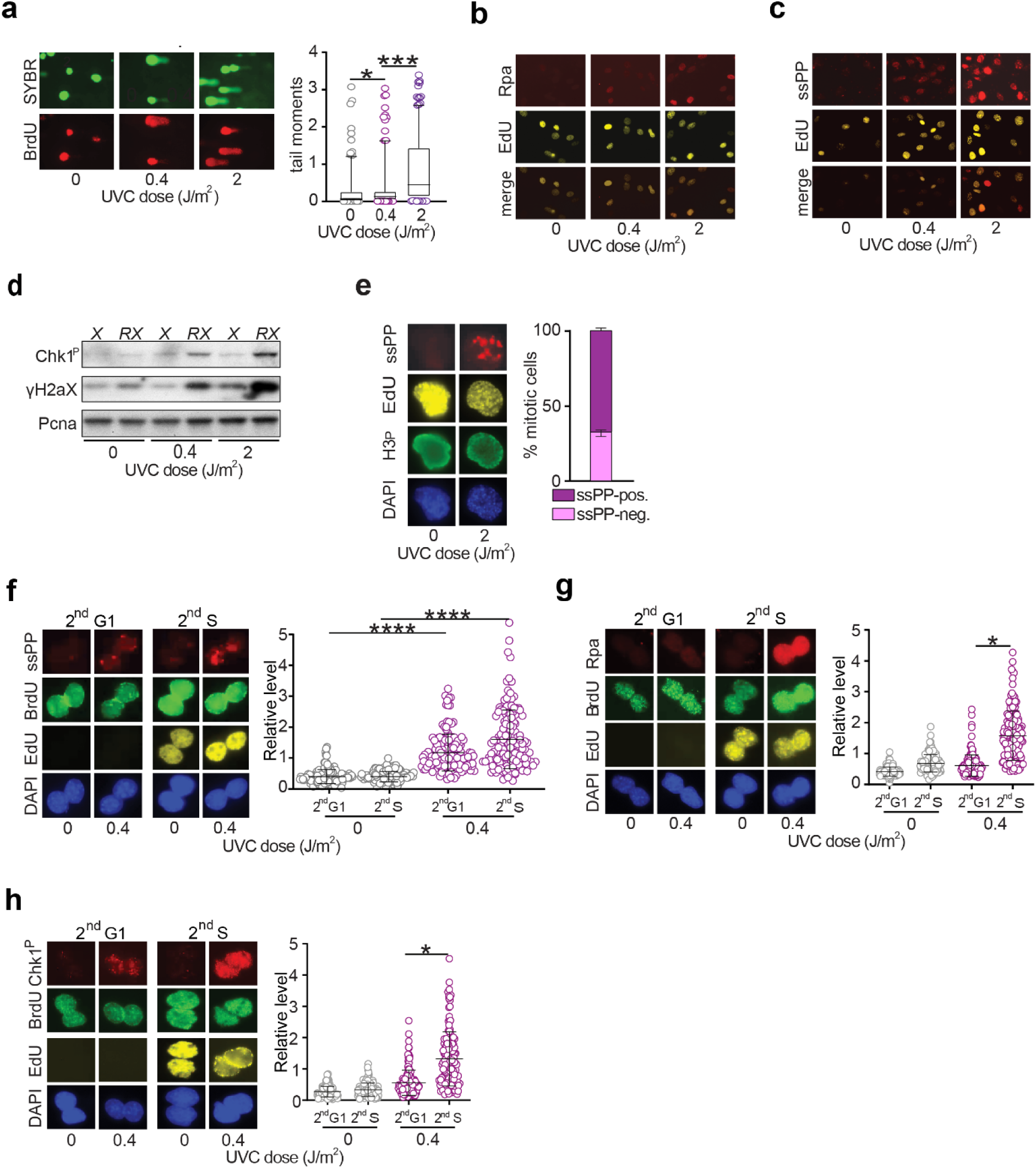
Induction of UV lesion-containing ssDNA tracts that are transmitted through mitosis into the next cell cycle. a) Left panel: alkaline comet electrophoresis to detect ssDNA breaks at 16 hours after low-dose UVC exposure of *Rev1Xpc* cells during S phase (BrdU^+^ cells). SYBR green was used to visualize DNA. Cells were arrested at mitosis with Nocodazole. Right panel: quantification of tail moments of BrdU^+^ cells from three different alkaline comet experiments. The box-and-whisker plots represent 5 to 95 percentiles. *: p<0.05, ***: p<0.001 (Student’s T test). b) Chromatin-associated Rpa at 8 hours after UVC exposure of S phase (EdU^+^) cells. c) ss(6-4)PP (ssPP) at 8 hours after UVC exposure of S phase (EdU^+^) cells. d) Western blot of DNA replication stress markers phosphoryated Chk1 (Chk1^p^) and γH2ax at 8 hours after UVC exposure of *Xpc* (*X*) and *Rev1Xpc* (*RX*) cells. Pcna: loading control. Cells were arrested at mitosis with Nocodazole. e) Left panel: staining for ss(6-4)PP (ssPP) in *Rev1Xpc* cells at 16 hours after UVC exposure during S phase (EdU^+^) and culture in the presence of Nocodazole. Phosphorylated Histone H3 (H3^p^) serves as a marker for mitotic chromatin. Right panel: fraction of ss(6-4)PP-positive staining in EdU^+^ H3^p^ *Rev1Xpc* cells. N=3, error bars: S.D. f) Left panel: immunostaining for ss(6-4)PP (ssPP) in binucleated G1 and S phase *Rev1Xpc* cells. Right panel: quantification of immunostaining of ss(6-4)PP in binucleated G1 and S-phase *Rev1Xpc* cells. Error bars: S.D. ****: p<0.0001 (Student’s T test). g) Left panel: immunostaining for ssDNA-bound Rpa in binucleated G1 and S phase *Rev1Xpc* cells. Right panel: quantification of ssDNA-bound Rpa in binucleated G1 and S phase *Rev1Xpc* cells. Error bars: S.D. *: p<0.05 (Student’s T test). h) Left panel: immunostaining for phosphorylated Chk1 (Chk1^p^) in binucleated G1 and S phase *Rev1Xpc* cells. Right panel: quantification of phosphorylated Chk1 in binucleated G1 and S phase *Rev1Xpc* cells. Error bars: S.D. *: p<0.05 (Student’s T test).

### Cells carrying unreplicated photolesions traverse the second cell cycle

Under-replicated fragile sites ^44^, Parp inhibitor–induced ssDNA tracts ^45^, and mismatch repair (MMR)–associated ssDNA tracts ^46, 47^ are known to persist through mitosis into the second cell cycle. To determine whether unreplicated 6-4)PP lesions similarly progress into the second cell cycle, we treated cells as described in Fig. 4a and stained these for ss(6-4)PP. Indeed, ss(6-4)PP lesions were still present in both G1 (BrdU⁺EdU⁻) and S (BrdU⁺EdU⁺) phases of the second cycle (Fig. 5f). This demonstrates that unreplicated (6-4)PP can traverse mitosis and persist to, at least, the subsequent S phase. Importantly, these persistent ss(6-4)PP lesions coincide with the formation of recombinogenic delayed DSBs during the second S phase (Fig. 4b–4e).

To assess DNA damage signaling during the second cell cycle, we stained *Rev1Xpc* cells for chromatin-bound Rpa and for Atr-phosphorylated Chk1. Both markers were significantly elevated during the second S phase (Fig. 5g, 5h), likely reflecting processing of delayed DSBs into 3′ ssDNA overhangs coated by Rpa, which amplifies Atr signaling ^12^. The dependence of delayed DSB induction on both persistent ss(6-4)PP and second-cycle replication suggests that these breaks arise from replication runoff or replisome disassembly at gapped templates opposing ss(6-4)PP.

### Delayed UVC-induced clastogenesis operates in wild type cells

To exclude the possibility that delayed clastogenesis is a particularity of *Rev1Xpc* and *Xpc* cells, we examined the kinetics of DSB formation in wild-type MEFs. Similar to mutant cells, wild-type MEFs exhibited a rapid increase in phosphorylated pKap1 shortly after UVC exposure (Fig. 6a), indicating that subtoxic (Fig. S1) UVC doses induce early DSBs also in wild type cells. At 24 hours post-exposure, only a few micronuclei were detected in binucleated wild-type cells (Fig. 6b) suggesting that, like in the mutant cells, most of the early DSBs were repaired before entry into the second cell cycle. However, when Cytochalasin B was added only at 24 hours, rather than immediately after exposure, the micronuclei frequency had increased markedly at 48 hours (Fig. 6b), consistent with a delayed mechanism of DSB induction similar to that observed in *Rev1Xpc* and *Xpc* cells.

**Figure 6:**
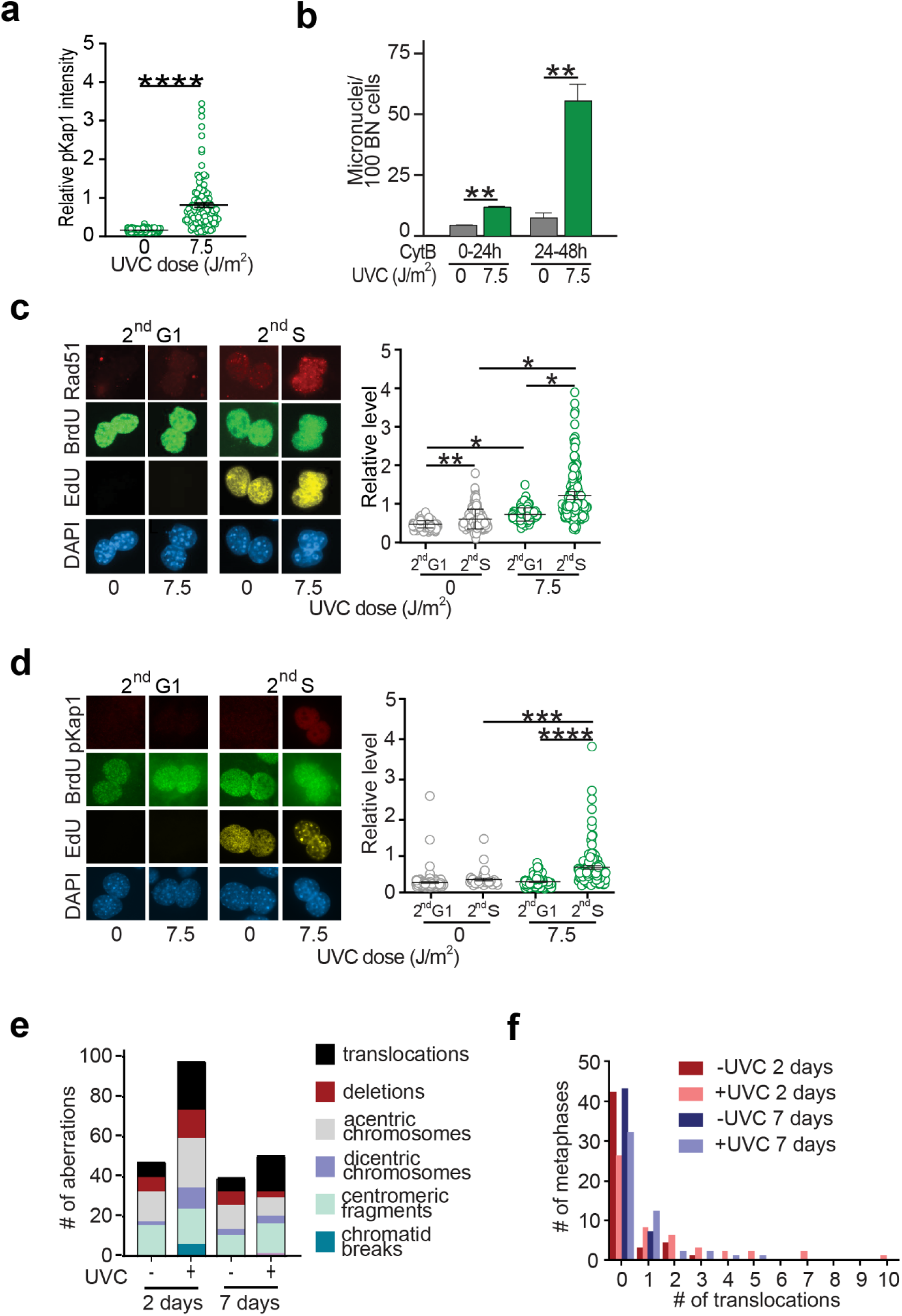
Delayed clastogenesis in wild type cells exposed to a subtoxic dose of UVC light. a) Quantification of immunostainings for phosphorylated Kap1 (pKap1) in EdU positive wild type cells at 0 and 4 hours after UVC exposure (7.5 J/m^2^). Cells were pulse-labelled with the replication marker EdU, immediately following exposure. Cells were arrested at mitosis using Nocodazole. Error bars: S.E.M. ****: p<0.0001 (Student’s T test) b) Quantification of micronuclei in binucleated (BN) wild type cells at 24 hours post-exposure and addition of Cytochalasin B (CytB) (0-24h) or at 48 hours post-exposure when CytB was added 24 hours post-exposure (24-48h). Error bars: S.E.M. **: p<0.01 (Student’s T test) c) Left panel: immunostainings for chromatin-bound Rad51 in binucleated G1 and S phase wild type cells. Right panel: quantification of chromatin-bound Rad51 in second cycle G1 and S phase wild type cells. Error bars: S.D. *: p<0.05, **: p<0.01 (Student’s T test). d) Left panel: immunostainings for pKap1 in binucleated G1 and S phase wild type cells. Right panel: quantification of pKap1 in second cycle G1 and S phase wild type cells. Error bars: S.D. ***: p<0.001, ****: p<0.0001 (Student’s T test). e) Characterization and quantification of chromosomal aberrations in 50 metaphases of wild type cells stained by COBRA-FISH, 2 or 7 days after mock treatment or exposure to 7.5 J/m^2^ of UVC light. f) Quantification of stable translocations per metaphase of wild type cells, 2 or 7 days after mock treatment or exposure to 7.5 J/m^2^ UVC light. Metaphases with 3 or more stable translocations are overrepresented at 2 (Pearson X^2^ test, p≤0.001) and at 7 days (Pearson Χ^2^ test, p≤0.05) after UVC exposure.

We next analyzed wild-type MEFs treated as in Fig. 4a for chromatin-bound Rad51 and pKap1 in second-cycle binucleated cells. Both markers were significantly elevated during the second S phase (BrdU⁺EdU⁺ cells) (Fig. 6c, 6d), mirroring the *Rev1Xpc* and *Xpc* cells. Thus, also wild-type cells generate DSBs in a delayed fashion following subtoxic UVC exposure.

To assess the induction and persistence of stable clastogenic events in wild type cells, we performed chromosome-specific painting of metaphase spreads at 2 and 7 days post-UVC exposure. Two days after treatment, mitotic wild type cells displayed numerous unbalanced rearrangements, including acentric fragments and translocations (Fig. 6e, 6f; Supplementary Fig. 5), indicative of the induction of clastogenesis by error-prone DSB repair ^24^. By day 7, fewer aberrations were observed (Fig. 6e–6f; Supplementary Fig. 5), likely reflecting a proliferative disadvantage of cells harboring unbalanced rearrangements.

Collectively, these findings indicate that most clastogenic DSBs induced by low-dose UVC represent the collapse of ss(6-4)PP lesions during the second S phase after exposure, irrespective of the genotype of the cells. Repair of these delayed DSBs generates sister chromatid exchanges and structural rearrangements or, if transmitted through the second mitosis, micronuclei at the onset of the third cycle as well as stable rearrangements. Thus, clastogenicity of low-dose UVC requires at least two cell cycles.

## Discussion

Processive DNA replication can be impeded by multiple factors, including nucleotide pool imbalances, transcription complexes, secondary DNA structures, lesions or incorporated ribonucleotides in the template, replication dysregulation driven by oncogene activation, and lesions at the template ^7^. At high lesion densities, clastogenesis has traditionally been assumed to originate from DSBs at collapsed replication forks, followed by error-prone end joining ^11, 24, 48, 49, 50^. However, to our knowledge, no conclusive evidence exists for this mechanism of clastogenesis.

Here, we demonstrate that, at least at low lesion densities, clastogenesis does not originate from stalled and collapsed replication forks. Our data rather suggest that most early DSBs resulting from stalled and collapsed replication forks are repaired rapidly. In contrast, lesion-containing ssDNA tracts, stabilized by RPA, do persist through mitosis, and these collapse into clastogenic DSBs only during the S phase of the subsequent cell cycle. This conclusion is supported by several observations: (i) cells carrying unreplicated lesions progress through mitosis and into the second cycle, likely facilitated by checkpoint tolerance ^13^; (ii) DSB formation peaks during the second S phase; (iii) second-cycle mitoses exhibit extensive chromosomal reshuffling via error-prone end joining and sister chromatid exchanges via homologous recombination; (iv) micronuclei appear at the onset of the third cycle, presumably representing unrepaired DSBs transmitted through the second mitosis.

We found that, following low-dose UVC exposure, rare cells are permanently arrested in the exposure cycle carrying numerous DSBs (Fig. 6e). We hypothesize that such cells carry an excessive amount of photolesions and therefore might suffer from the consequence of Rpa exhaustion resulting in “early” clastogenesis that exacerbates replication fork collapse ^13^. However, our data show that delayed genome instability originating during the second S phase, represents a more physiological response to non-saturating lesion densities. Although evident in wild-type cells, this mechanism is exacerbated by TLS and NER defects, underscoring the roles of these pathways in suppressing clastogenesis.

Conversely, the enrichment of deleterious NER and HR mutations in highly rearranged prostate carcinomas (Fig. 1b, 1c) supports a causal link between persistent nucleotide lesions and cancer etiology. These findings suggest that delayed clastogenicity of unreplicated lesions may broadly contribute to genome instability in cancers associated with chronic exposure to DNA damage, and that this phenotype is exacerbated by somatic NER or HR defects in the cancer.

The transfer of ssDNA tracts through mitosis is not unprecedented ^44, 45, 46, 47^, but has not previously been linked to clastogenesis or carcinogenesis. Notably, mismatch repair (MMR)-induced excision tracts can persist through mitosis and collapse into DSBs during the next S phase ^46, 47^, predicting that MMR deficiency may protect against clastogenesis—a hypothesis supported by the scarcity of structural genome aberrations in MMR-deficient tumors ^51^.

Our results provide mechanistic insight into clastogenesis at low lesion densities and its relevance to genome-rearranged cancers. These findings also have implications for clastogenicity testing: the cytokinesis-block micronucleus assay ^52^ should incorporate delayed Cytochalasin B addition and monitor micronuclei only after the second mitosis. Finally, the *Rev1Xpc* cell line and assays described here may serve as a highly sensitive model for detecting clastogenic potential of DNA-damaging agents.

## Acknowledgements

This work was supported by a PhD scholarship to P.T. from the Office of the Higher Education Commission, the Royal Thai Government. A.T.-S. was funded by grant 11358 from the Dutch Cancer Society to N.d.W. D.C.dG was supported by a grant of the Dutch cancer foundation NKI-2017-10796 to H.J. We thank Kajal Ramkisoensing for technical assistance and Robbert Ijsselsteijn for help with statistical analyses. We thank the Hartwig Medical Foundation (https://www.hartwigmedicalfoundation.nl) for making sequences of prostate tumors available for analysis.

## Conflict of interested statement

The authors declare no conflicts of interest.

## Sharing of data and reagents

All primary data and unique reagents described will be made available upon motivated request.

## Methods

### Analysis of cancer genome sequences

The whole-genome sequencing (WGS) database of the Hartwig Medical Foundation was used to select data from patients with metastatic prostate cancer that were under the protocol of the Centre for Personalized Cancer Treatment (CPCT, <u>NCT01855477</u>). Detailed methods on sequencing protocols, SNP calling and structural variances analysis are as described ^53^. Proficiency or deficiency of pathways were decided upon whether a tumor had a predicted deleterious mutation according to SIFT ^54^ in one or more of the genes associated with the pathway (Supplementary table 2). Frameshift mutations were considered as deleterious as well. Interchromosomal rearrangements were defined as a fragment starting and ending at two different chromosomes according to the structural variance pipeline as described ^53^. Data was sorted and analyzed with R version 3.6.3, whereas statistical analyses was performed using Graphpad Prism7 software. P values where calculated using Fisher’s exact test (Supplementary table 2). Cutoff was selected based on frequency distribution of interchromosomal rearrangements of NER proficient and deficient tumors.

### Cell culture

The generation of isogenic MEF lines, wild type or deficient for *Xpc*, *Rev1* or double-deficient for *Rev1* and *Xpc* has been described previously ^36^. Cells were cultured in Dulbecco’s modified Eagle’s medium (DMEM) containing 4.5 g/l glucose, Glutamax and pyruvate (Invitrogen), supplemented with 10% fetal calf serum, penicillin (100 U/ml), and streptomycin (100 μg/ml) (DMEM medium) at 37°C in a humidified atmosphere containing 5% CO_2_.

### Clonogenic survival assay

MEFs were treated with UVC doses of up to 10 J/m^2^, trypsinized, and counted. A fixed number of cells was plated in 9 cm dishes and cultured for 10 days at 5% CO_2_ and 37°C. Clones were counted after staining with methylene blue. To determine the relative clonogenic survival, the cloning efficiency of unexposed cells was set at 100%.

### Cytokinesis-blocked micronucleus assays

The assay was performed as described previously ^43, 55^. Briefly, 7.5 × 10^4^ cells were plated on a glass slide (76 mm × 26 mm) and cultured overnight. Prior to exposure, cells were washed twice with PBS. At 0 or 24 hours after UVC treatment, Cytochalasin B (3 μg/ml; Sigma-Aldrich) was added to the cultures in order to inhibit cytokinesis. Twenty-four hours after addition of Cytochalasin B, cells were fixed using 3.5% paraformaldehyde (4°C) and 0.5% Triton-X100. The slides were washed trice with PBS. Subsequently, the slides were soaked in 70%, 90% and absolute ethanol for 5 min each, respectively. Then, slides were rinsed with PBS and nuclei were stained with 4′,6-diamidino-2-phenylindole (DAPI, 17.5 ng/ml). The binucleated cells and micronuclei were scored using a fluorescence microscope (Zeiss, Germany) and the Metafer 4 program (Metasystem, Germany). Three independent experiments were performed and mock-treated cells were included as a control.

To determine responses in the second cell cycle following UVC treatment, cells were incubated with 10 μM Bromodeoxyuridine (BrdU, Millipore) for 30 min after which the cells were cultured in the presence of Cytochalasin B for 23 hours to generate binucleated cells. Micronuclei in binucleated cells showing BrdU-positive nuclei are derived from S phase cells at the time of UV exposure (first cell cycle micronuclei). To distinguish binucleated cells with nuclei in G1 phase from those that have entered S phase, the cells were incubated in medium containing 10 μM 5-ethynyl-2’-deoxyuridine (EdU, Invitrogen) for 30 min, prior to fixation. Consequently, binucleated cells with nuclei double-positive for BrdU and EdU reflect 2^nd^ S phase cells after UVC exposure. BrdU in DNA was visualized after depurination of nuclear DNA with 2 M HCl (30 min.). Before incubation with anti-BrdU antibodies, coverslips were rinsed intensively to quench the acidity. BrdU was visualized using an Alexafluor 555-labeled goat-anti-mouse antibody according to the manufacturer’s protocol. EdU in DNA was visualized using Alexafluor 647-conjugated azide (Click-iT^TM^ EdU imaging kit, Invitrogen) according to the manufacturer’s recommendation.

### Analysis of genome rearrangements by multicolor COBRA-FISH karyotyping

Cells were exposed to 7.5 J/m^2^ UVC and cultured for 2 or 7 days. The metaphase harvest, FISH and karyotyping procedures were performed as described ^56^.

### Bivariate cell cycle analysis

MEFs were seeded in 90 mm dishes and were cultured overnight to 70% confluence. Cells were rinsed with PBS and exposed to UVC or mock-treated. Immediately after exposure to 0-2 J/m^2^ UVC, cells were pulse labelled with BrdU for 30 min in 5% CO_2_ at 37°C. Cells were trypsinized and fixed with 70% ethanol at indicated time points. BrdU staining and flow cytometry were performed as described ^43^.

### Single-cell gel electrophoresis (comet) assays

DNA strand interruptions and dsDNA breaks in cells that were replicating when exposed to UVC were measured by alkaline and neutral comet assays, respectively. In detail, cells were cultured at 60-70% confluence and pulse-labeled for 30 min with BrdU, immediately after exposure to UVC. This enables the identification of replicating cells at the time of UVC treatment. Then, the cells were cultured in the presence of nocodazole (300 ng/ml) for 16 hours to accumulate cells at mitosis. Single-cell suspensions were processed to visualize comets according to the manufacturer’s instructions (Trevigen). After gel electrophoresis, chromatin was depurinated and stained for incorporated BrdU as above. DNA was visualized using SYBR green (Invitrogen). The comet tail moments of 180-240 BrdU-containing nuclei from three independent experiments were scored using Comet analysis software (TriTek).

### Immunofluorescence

The antibodies used in immunofluorescence were mouse anti-BrdU (Beckton Dickinson), rabbit anti-BrdU (Rockland immunochemicals), mouse anti-Histone H3^S10-P^ (Millipore), rabbit anti-Histone H3^S10-P^ (Millipore), mouse anti-(6-4)PP (CosmoBio), rabbit anti-Chk1^S345-P^ (Cell signaling), rat anti-Rpa (Cell signaling), mouse anti-Rad51 (Gentex), rabbit anti-Kap1^S824-P^ (Bethyl) and appropriate secondary antibodies conjugated with fluorescence dyes (Invitrogen). To detect ss(6-4)PPs, Rpa and Chk1^S345-P^ in the late S/G2 phase, cells were seeded on coverslips and cultured overnight in 5% CO_2_ at 37°C. Prior to exposure to 0–2 J/m^2^ UVC, cells were pulse-labeled with EdU for 30 min. After UVC exposure, cells were cultured in the presence of nocodazole for 8 hours, to prevent cells from entering a second cell cycle after exposure. Then cells were fixed and immunostained for ss(6-4)PPs ^36^, Rpa or Chk1^S345-P^, and for the mitotic marker H3^S10-P^, using appropriate antibodies. To increase the sensitivity of anti-(6-4)PPs staining, Tyramine Signal Amplification (TSA, Perkin-Elmer) was applied, according to the manufacturer’s recommendations. Late S/G2 phase cells that were replicating at the time of UV exposure were identified by positive staining for EdU and negative staining for H3^S10-P^. To detect ss(6-4)PPs in mitotic cells, cultures were treated as described above, with the exception that the cells were cultured in nocodazole-containing medium for 16 hours to accumulate G2/M phase cells. The mitotic cells that were replicating at the time of UVC treatment were identified by positive staining for both EdU and H3^S10-P^. DNA damage response proteins and ss(6-4)PPs in binucleated cells were visualized as follows. Cells were cultured on coverslips in medium with 10 μM BrdU for 30 min, immediately after irradiation with 0-7.5 J/m^2^ UVC. Then, cells were cultured in the presence of Cytochalasin B for 16 hours. Prior to fixation, cells were pulse-labeled with 10 μM EdU in the presence of Cytochalasin B for 30 min. Since BrdU detection involves a depurination step, which may destroy protein epitopes, fixed cells were firstly stained for EdU. Then, cells were blocked with 5% BSA+0.1% tween-20 in PBS for 30 min and subsequently incubated overnight with anti-Rpa, Chk1^S345-P^, Rad51 or Kap1^S824-P^ antibodies diluted in 1% BSA+0.1% tween-20 in PBS at 4^0^C. Then, appropriate secondary antibodies were applied. After washing, cells were again fixed with 3.7% paraformaldehyde for 15 min, followed by incubation with 2N HCl for 12 min. After incubation with anti-BrdU antibodies, an appropriate secondary antibody was applied and nuclei were stained with DAPI. ss(6-4)PPs were visualized as described above. Images were captured using a wide-field microscope (Zeiss Axioplan 2 or M2). Image intensities of at least 150 binucleated cells per condition from three independent experiments were quantified using Fiji (ImageJ, National Institutes of Health).

### Western blotting

For Western blotting, cells were exposed to 0, 0.4 or 2 J/m^2^ UVC light and cultured in the presence of nocodazole for 8 hours. Cells were lysed by adding Laemmli lysis buffer. Proteins were separated by SDS-PAGE and blotted onto a nitrocellulose membrane (Protran, Amersham Biosciences). After incubation in blocking buffer [(Rockland) and PBS-0.1% Tween 20 (Sigma-Aldrich)] for at least 1 hour at room temperature, the membrane was incubated overnight with rabbit monoclonal anti-phospho Chk1 (S345) (1:2000; Cell Signaling), mouse monoclonal anti-phospho-H2ax (S139) (1:4000; Millipore) and mouse monoclonal anti-PCNA antibody (1:8000; Santa Cruz) at 4°C. Proteins were visualized by enhanced chemiluminescence detection after incubation with appropriate peroxidase-conjugated secondary antibodies (Bio-Rad) for 45 min at room temperature.

### Analysis of mitotic aberrations

MEFs were exposed to UVC and immediately labelled with 10 μM BrdU for 22 hours. To analyze mitoses of the second cell cycle that follows exposure during S phase, MEFs were exposed to UVC and labeled with BrdU for 22 hours. Then, BrdU-containing medium was removed and cells were labelled with 10 μM EdU for 18 hours. At 15 hours of EdU labelling, 10 μg/ml Colcemid was added for 3 hours. Then, cells were fixed and stained for BrdU and EdU. BrdU^+^, EdU^+^ metaphase spreads were analyzed for the distribution of BrdU and EdU labelling over sister-chromatids and for aberrations. The experiment was performed three times. Quantification of aberrant chromosomes was performed on 15–31 metaphases per experiment.

