## Supplementary Figures and Tables for "Clastogenesis by nucleotide lesions requires the completion of two cell cycles"

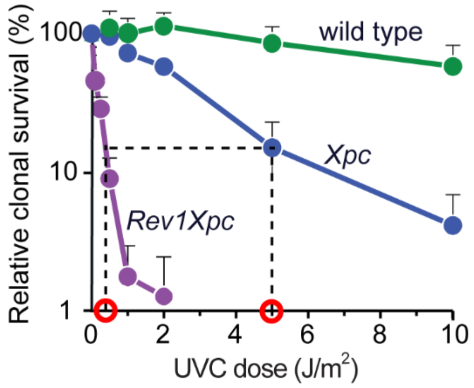


Figure S1

**Supplementary figure S1: Sensitivity of MEF lines for UVC light.**

Relative clonal survival of MEF lines in response to UVC light exposure. Red circles: equitoxic doses. N=3. Error bars: S.D.

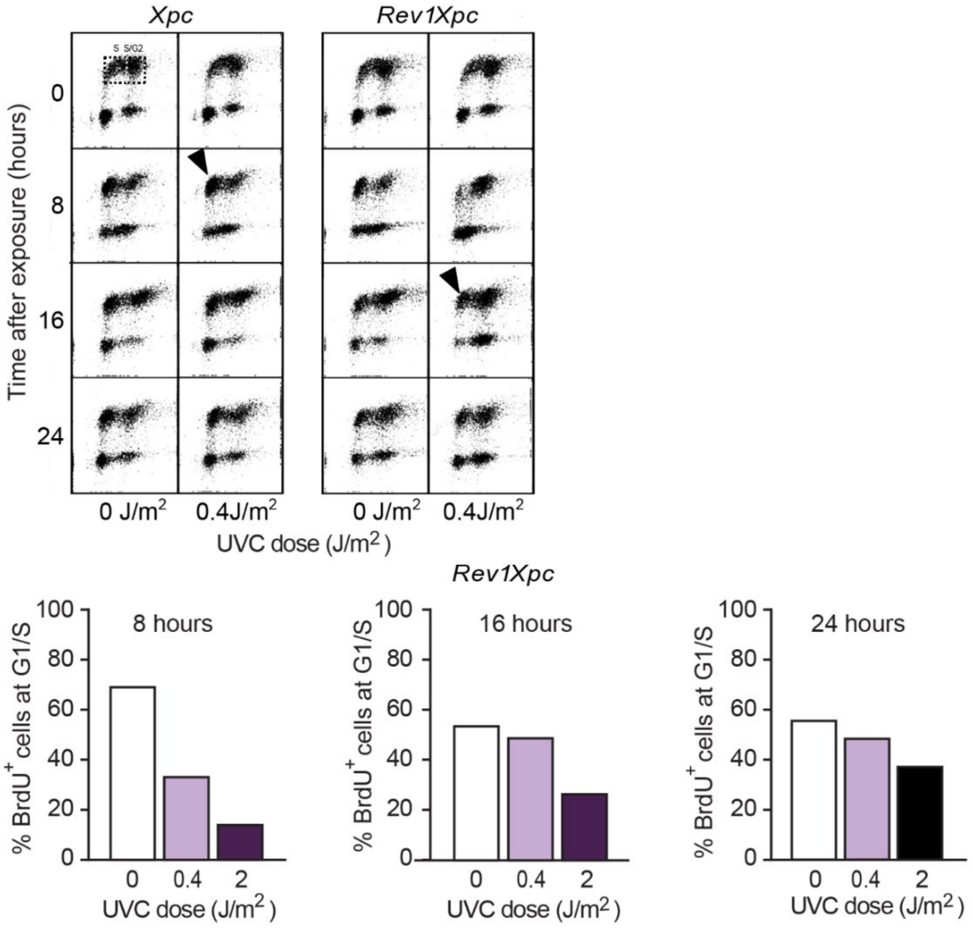


\

Figure S2

**Supplementary figure S2: Cell cycle progression of *Xpc* and *Rev1Xpc* cells following low dose UVC exposure.**

Upper panel: FACS plots of cells that were pulse-labeled with BrdU immediately after mock treatment or exposure to 0.4 J/$m^2$ UVC and cultured for up to 24 hours after treatment. Horizontal axis: DNA content per cell, Y axis: incorporated BrdU. Arrows display progression of BrdU positive cells to the next cell cycle. *Rev1Xpc* cells show delayed progression to the second cell cycle following 0.4J/$m^2$ UV exposure. Lower panel: quantification of $BrdU^+$ *Rev1Xpc* cells that reappear at the G1/early S phases of the next cell cycle following mock treatment or exposure to 0.4 – 2 J/$m^2$ of UVC. Cells were pulse-labelled with BrdU immediately after UV exposure or mock treatment.

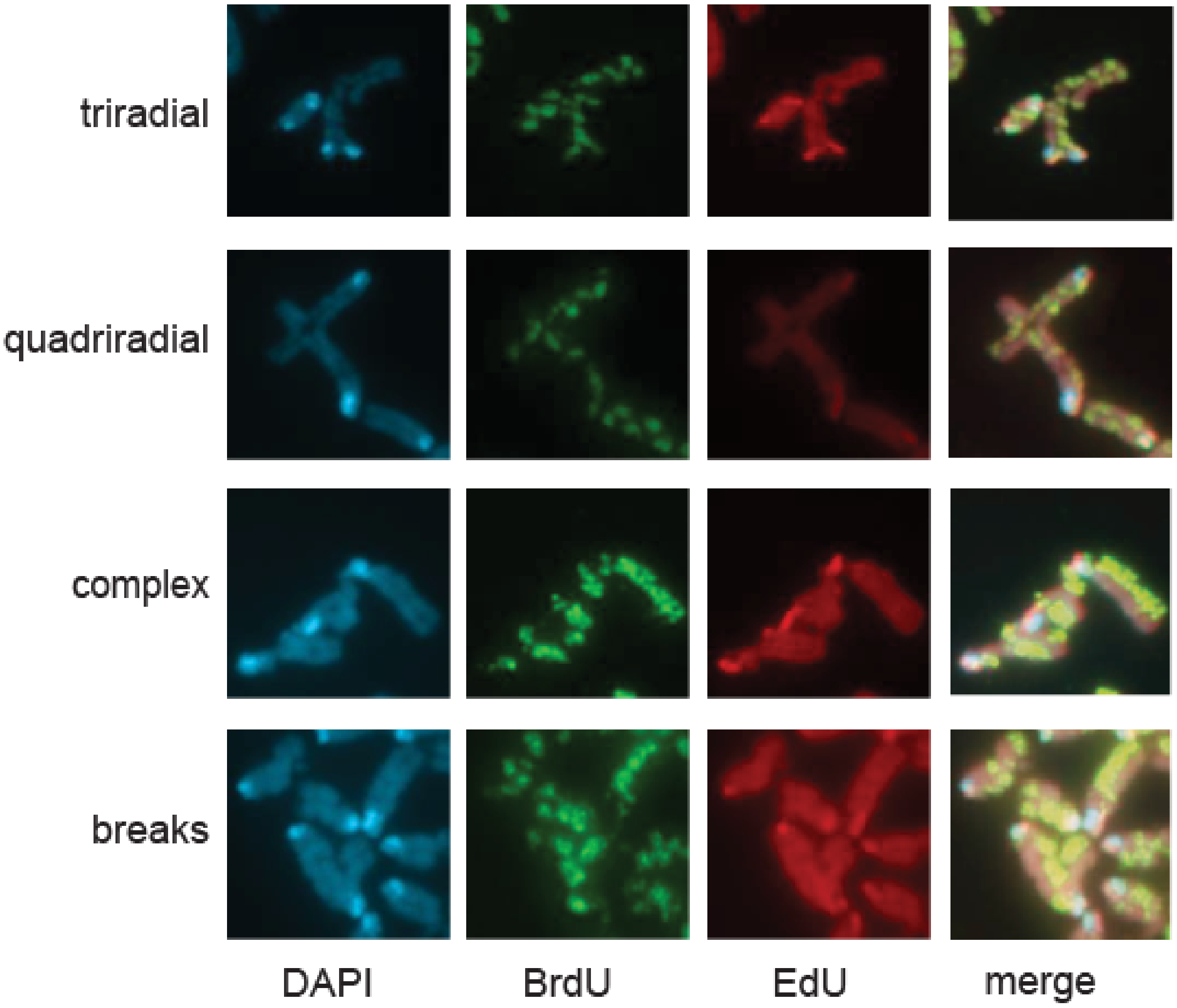


Figure S3

**Supplementary figure S3: Aberrant chromosomes following UVC exposure.**

*Rev1Xpc* MEFs were labelled with BrdU (S phase of exposure) immediately upon UVC exposure (0,4 J/$m^2$), and with EdU (second S phase), at 22 hours after exposure. Blue: DAPI. Cells were arrested at 40 hours beyond exposure using Colcemid. These paired chromatids are believed to be intermediates in the generation of chromosomal rearrangements. White arrow: chromatid break.

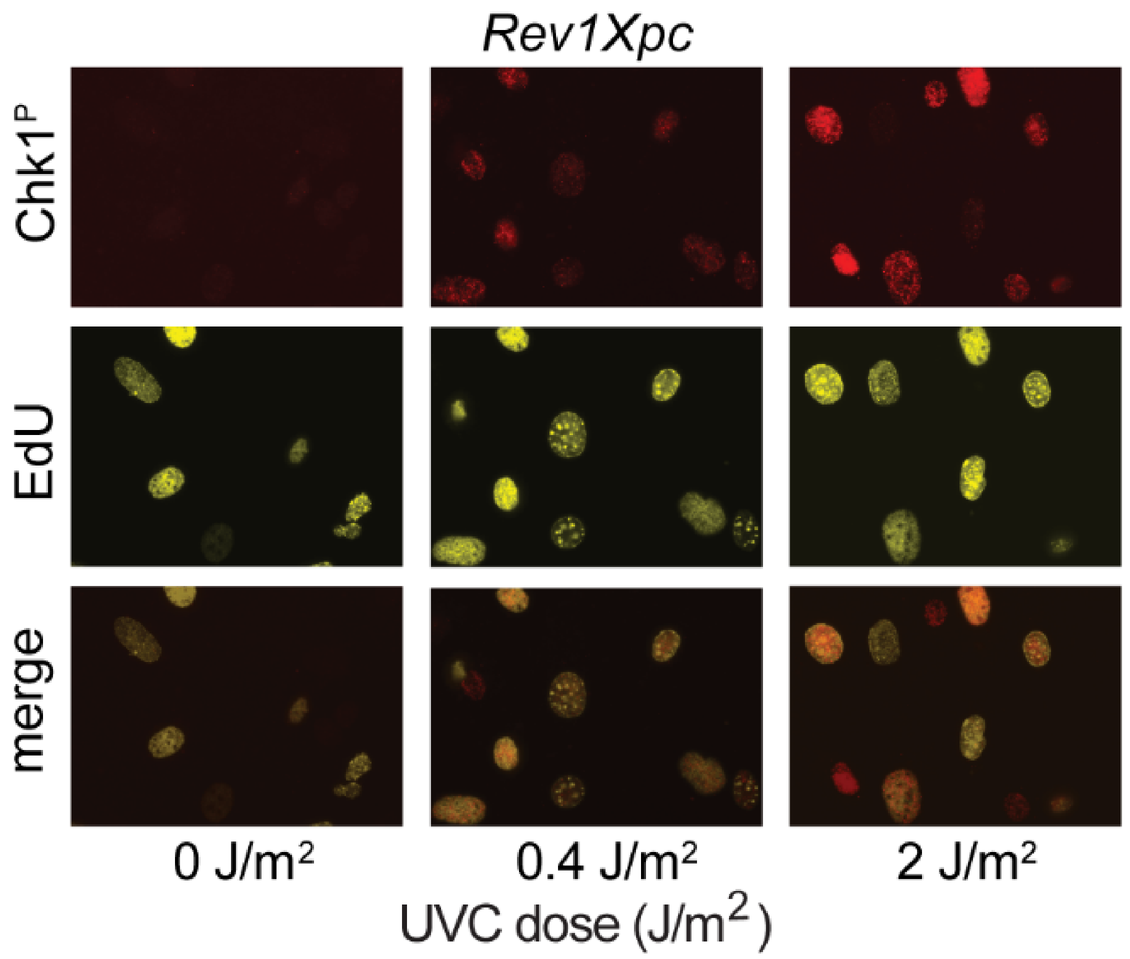


Figure S4

**Supplementary figure S4: Chk1 phosphorylation in *Rev1Xpc* cells following low dose UVC exposure.**

Immediately following UVC exposure, S phase *Rev1Xpc* cells were pulse-labelled with EdU. Eight hours after UVC exposure, cells were fixed and (immuno)stained for EdU and for phosphorylated Chk1 ($Chk1^P$).

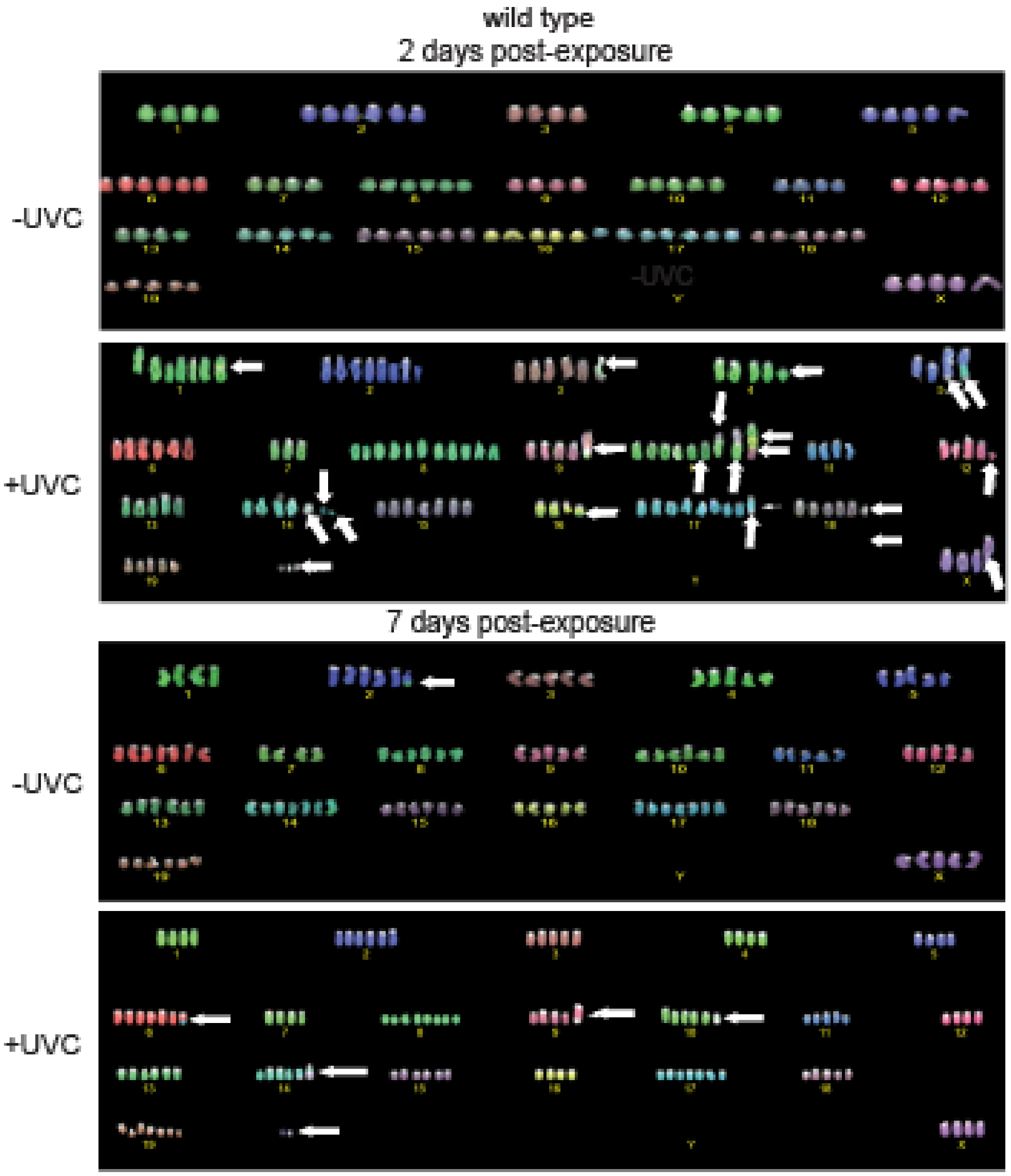


Figure S5

**Supplementary figure S5: Chromosomal aberrations in wild type cells visualized by COBRA-FISH.**

COBRA-FISH analysis of wild type MEFs, 2 or 7 days after mock treatment or exposure to a mildly toxic UVC dose (7.5 J/m$^2$) showing stable chromosomal aberrations (white arrows).

**Supplementary table 1: Genes analyzed for the presence of mutations in prostate carcinomas.**

| Pathway | Genes |
|---|---|
| **NER** | *Ccnh, Cdk7, Cetn2, Ddb1, Ddb2, Ercc1, Ercc2, Ercc3, Ercc4, Ercc5, Ercc6, Ercc8, Gtf2h1, Gtf2h2, Gtf2h3, Gtf2h4, Gtf2h5, Lig1, Mms19, Mnat1, Rad23a, Rad23b, Rpa1, Rpa2, Rpa3, TfIIh, Uvssa, Xab2, Xpa, Xpc* |
| **HR** | *Brca1, Brca2, Rad51, Rad51b, Rad52, Xrcc2, Xrcc3* |
| **FA** | ***Atr***, ***Atrip***, *Blm, Brca1, Brca2, Brip1, Eme1, Eme2, Ercc1, Ercc4, Faap24, Faap100, Fan1,* ***Fanca, Fancb, Fancc, Fancd2, Fance, Fancf, Fancg, Fanci, Fancl, Fancm***, *Hes1, Mlh1, Mus81, Palb2, Pms2,* ***Polh, Poli, Polk, Poln,*** *Rad51,* ***Rev1, Rev3l***, *Rmi1, Rmi2, Rpa3, Rpa4, Slx4, Telo2, Ube2t,* ***Usp1,*** *Wdr48* |

NER: nucleotide excision repair.

HR: double-strand breaks repair by homologous recombination.

FA: crosslink repair by the Fanconi Anemia pathway. Overlaps with TLS-associated genes (in bold).

**Supplementary Table 2. Predicted deleterious somatic mutations in genes associated with NER, HR and FA, in genome-instable (>200 rearrangements) prostate carcinomas.**

| | Over 200 rearrangements (n=15) | Up to 200 rearrangements (n=339) |
|---|---|---|
| **NER-** | 4 | 22 |
| **NER+** | 11 | 317 |
| **HR-** | 4 | 26 |
| **HR+** | 11 | 313 |
| **FA-** | 5 | 69 |
| **FA+** | 10 | 270 |

NER: nucleotide excision repair.

HR: double-strand breaks repair by homologous recombination.

FA: crosslink repair by the Fanconi Anemia pathway. Overlaps with TLS-associated genes.
